# iMDA-BN: Identification of miRNA-Disease Associations based on the Biological Network and Graph Embedding Algorithm

**DOI:** 10.1101/2020.07.01.181982

**Authors:** Kai Zheng, Zhu-Hong You, Lei Wang

## Abstract

Benefiting from advances in high-throughput experimental techniques, important regulatory roles of miRNAs, lncRNAs, and proteins, as well as biological property information, are gradually being complemented. As the key data support to promote biomedical research, domain knowledge such as intermolecular relationships that are increasingly revealed by molecular genome-wide analysis is often used to guide the discovery of potential associations. However, the method of performing network representation learning from the perspective of the global biological network is scarce. These methods cover a very limited type of molecular associations and are therefore not suitable for more comprehensive analysis of molecular network representation information. In this study, we propose a computational model based on the Biological network for predicting potential associations between miRNAs and diseases called iMDA-BN. The iMDA-BN has three significant advantages: I) It uses a new method to describe disease and miRNA characteristics which analyzes node representation information for disease and miRNA from the perspective of biological networks. II) It can predict unproven associations even if miRNAs and diseases do not appear in the biological network. III) Accurate description of miRNA characteristics from biological properties based on high-throughput sequence information. The iMDA-BN predictor achieves an AUC of 0.9145 and an accuracy of 84.49% on the miRNA-disease association baseline dataset, and it can also achieve an AUC of 0.8765 and an accuracy of 80.96% when predicting unknown diseases and miRNAs in the biological network. Compared to existing miRNA-disease association prediction methods, iMDA-BN has higher accuracy and the advantage of predicting unknown associations. In addition, 45, 49, and 49 of the top 50 miRNA-disease associations with the highest predicted scores were confirmed in the case studies, respectively.

## 1 Introduction

MicroRNAs (miRNAs) are small, non-coding RNA molecules that affect basic biological processes by base pairing with targeted mRNAs [1, 2]. In particular, many studies have revealed that miRNAs act as negative gene regulators in a variety of human diseases and are involved in disease processes such as breast cancer, myasthenia gravis, primary biliary cirrhosis, and the like [3–5]. This suggests that miRNAs can promote the development of new therapeutic strategies by acting as biomarkers for certain diseases. Therefore, exploring how to predict the relationship between miRNA and disease on a large scale has always been a research hotspot in the field of bioinformatics.

In recent years, many predictive tools have been proposed that convert each node in the network (including miRNAs and diseases) into low-dimensional potential representations to calculate network representation associations in order to identify miRNA-diseases association in the context of known network structures. However, since most predictive tools only introduce intermediary to build a two-layer network, the amount of information represented by the network describing the nodes is relatively rare. For example, Shi *et al*. proposed a computational model for predicting potential miRNA-disease associations by integrating miRNA-target networks, gene-disease networks, and protein-protein interaction networks. This method introduces many networks but does not quantify the network representation information of nodes from the entire network [6]. Similarly, Mork *et al*. constructed a miRNA-protein-disease network for association prediction which contributed greatly to inferring potential associations from the network structure but was still not comprehensive enough [7]. Later, Yang *et al*. calculated miRNA functional similarity by constructing a miRNA gene network to improve the performance of miRNA-disease association prediction. The contribution of this method to the field is obvious, but the constructed two-layer network does not truly reflect the relationship between nodes in reality [8]. In addition, Chen *et al*. proposed a prediction model based on binary network projection called BNPMDA, which introduces Medical Subject Headings to describe disease information [9]. Furthermore, there are many predictive tools that use domain knowledge as a supplement to high-throughput data to improve prediction accuracy, such as gene ontology (GO), medical subject terms, and miRNA-disease-associated network information [10–12]. For example, lan *et al*. proposed a computational framework called KBMF-MDI, which uses miRNA sequence similarity to improve model performance [13]. Later, wang *et al*. used natural language processing techniques to extract miRNA sequence features for the first time in the miRNA-disease association prediction model, which made an important contribution to accurately describe miRNA characteristics [14].

In this study, we propose a novel miRNA-disease association predictor based on biological network that attempts to overcome the above problem, called iMDA-BN. Different from the previous method, the new predictor has the following characteristics: I) Biological networks composed of miRNA, lncRNA, protein, drug, and disease can describe the network representation of miRNAs and diseases from the perspective of the entire network, rather than being restricted to intermediaries. II) The association of pairs of new diseases and new miRNAs can be predicted and has a high degree of accuracy. III) Sequence information based on high-throughput sequencing is used to accurately quantify miRNA function information. In iMDA-BN, 9 relationship types are integrated, including miRNA-lncRNA, miRNA-disease, miRNA-protein, lncRNA-disease, lncRNA-protein, protein-disease, drug-protein, drug-disease, protein-protein and 105,546 associations to build the biological network to assist in predicting potential associations between miRNA and disease. To our knowledge, this is the largest biological network for predicting miRNA-disease associations. To verify the performance of the iMDA-BN, we applied it to the benchmark data set to achieve an AUC of 0.9145 with an accuracy of 84.49%. And when predicting the associations between new diseases and new miRNAs, it can achieve an AUC of 0.8765 with an accuracy of 80.96%. In addition, in order to verify the robustness of the proposed predictor, three diseases were used for case studies. From the performance in various evaluations, the proposed prediction model based on Biological Network can be used as a good tool for predicting tasks.

## 2 Materials and Methods

### 2.1 Data sets

#### 2.1.1 The baseline data set of miRNA-disease association

With the deepening of biomedical research, the demand for integrated miRNA disease association databases is also growing, and various public databases and benchmark data sets have emerged. They manually collected a large number of miRNA-disease association entries from the literature, for example, HMDD v3. 0, dbDEMC v2.0 and miR2Disease [15–17]. In this study, HMDD v3.0 was selected as the baseline database because it has the most comprehensive miRNA-disease association to date, collecting 32281 experimentally supported miRNA-disease associations consisting of 1102 miRNA genes and 850 diseases. Due to the update of the version and the overlap of evidence supporting the association, they were combined into a group of associations covering 1,206 miRNAs and 894 diseases. In this group, 901 miRNAs have sequence information in miRbase [18]. Therefore, the final data set included 16427 associations consisting of 901 miRNAs and 877 diseases were used in our experiments.

#### 2.1.2 The baseline data set of miRNA sequence information

With the development of high-throughput sequencing technology, biological characteristics such as miRNA sequence information have been gradually supplemented, and many public databases have begun to collect and integrate the biological information, including miRBase, PMRD and MicroRNAdb [19, 20]. Among them, miRbase has the most complete miRNA information, containing 24,521 microRNA loci from 206, which can process 30424 mature microRNA products. Therefore, in this experiment we downloaded high-throughput data from miRNAs from miRbase to complement miRNA sequence information. The database is accessible free of charge via the web server http://www.mirbase.org/.

#### 2.1.3 The Biological Network

The complex homogeneous network constructed with associations between molecules can use the network representation information of its nodes as features to predict associations. The Biological Network consists of known molecules, as shown in Figure 1. However, predictors based on the entire molecular network are rarely proposed. The biological network (BN) constructed by Guo *et al*. provides a new perspective. In order to fill this part of the research gap, we introduce the MAN into the prediction of miRNA-disease association. So far, the biological network consists of small biomolecule transcripts (proteins, lncRNAs and miRNAs), drugs and diseases. As shown in Table 1, there are nine kinds of associations which are miRNA-disease (MDA), miRNA-lncRNA (LMA), miRNA-protein (MPI), lncRNA-disease (LDA), lncRNA-protein (LPI), protein-disease (PDA), protein-protein (PPI), drug-protein (DPI), drug-disease (DDI).

**Figure 1.**
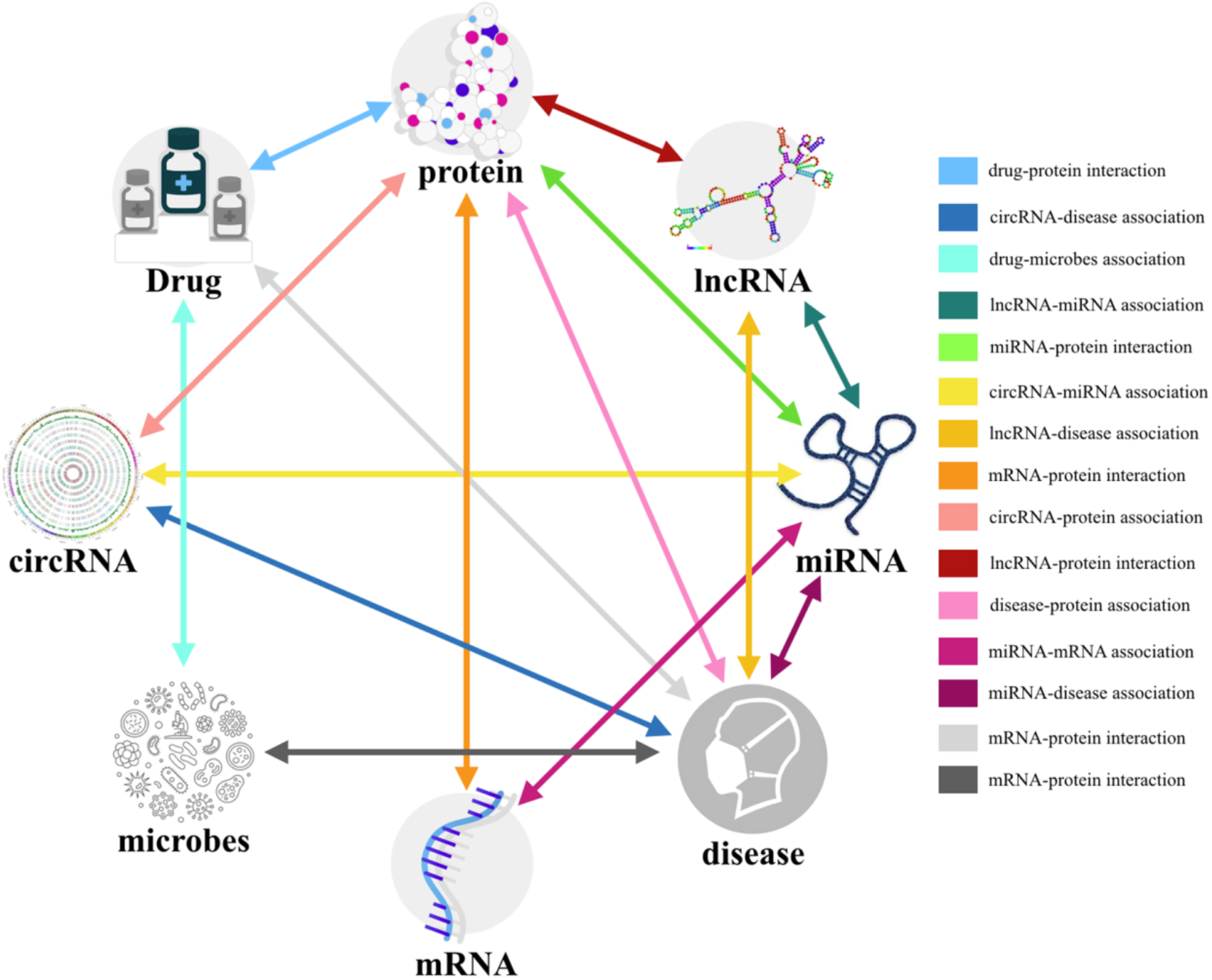
The biological network.

**Table 1.**
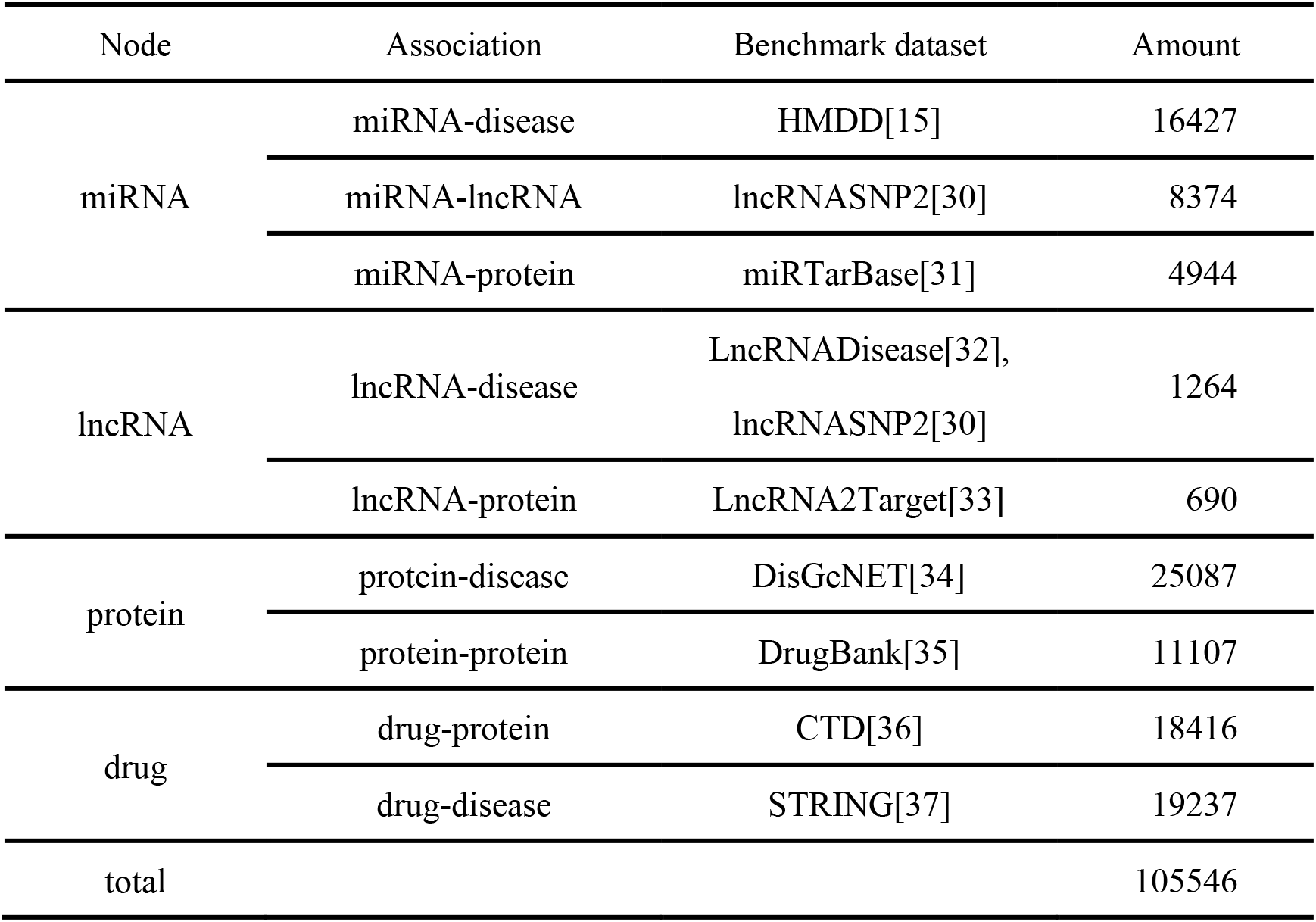
The nine associations that constitute the biological network.

| Node | Association | Benchmark dataset | Amount |
| --- | --- | --- | --- |
| miRNA | miRNA-disease | HMDD[15] | 16427 |
|  | miRNA-lncRNA | lncRNASNP2[30] | 8374 |
|  | miRNA-protein | miRTarBase[31] | 4944 |
| lncRNA | lncRNA-disease | LncRNADisease[32],<br>lncRNASNP2[30] | 1264 |
|  | lncRNA-protein | LncRNA2Target[33] | 690 |
| protein | protein-disease | DisGeNET[34] | 25087 |
|  | protein-protein | DrugBank[35] | 11107 |
| drug | drug-protein | CTD[36] | 18416 |
|  | drug-disease | STRING[37] | 19237 |
| total |  |  | 105546 |

Based on the above reference database, the number of various types of nodes in the statistical biological network is as shown in Table 2.

**Table 2.** The number of five nodes in the biological network.

| Node | MiRNA | LncRNA | Protein | Drug | Disease | Total |
| --- | --- | --- | --- | --- | --- | --- |
| Amount | 1023 | 769 | 1649 | 1025 | 2062 | 6528 |

### 2.2 Methods

#### 2.2.1 Attribute information of miRNAs and diseases

##### Semantic descriptor of disease

Disease descriptors describe disease attributes in medical subject vocabulary terms and organize them in Directed Acyclic Graphs (DAGs) where edges represent the association between diseases and nodes indicate disease. One of the core issues in the extraction of disease information is how to measure disease semantic relevance through the Medical Subject Headings (MeSH) terms [11]. In disease DAG, the association between the two diseases depends on their location/depth in the DAG, and if the two diseases have semantic similarities they will share more DAG parts. The semantic similarity S_sem_(*d*(*i*), *d*(*j*)), of disease *d*(*i*) and disease *d*(*j*) are defined as follows:

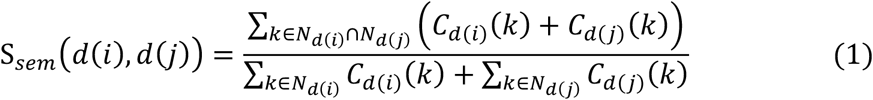

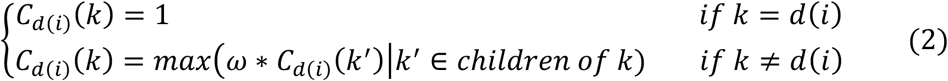

where *C*_*d*(*i*)_(*k*) is the semantic contribution of disease *k* to disease *d*(*i*). *ω* is the contribution coefficient, which is set to 0.5 according to the previous study[21]. *N*_*d*(*i*)_ is a collection of all diseases that appear in the DAG of disease *d*(*i*). When the semantic similarity between all diseases in the biological network is calculated, the semantic similarity matrix *S_sem_* whose size is 2062 × 2062 can be obtained. Therefore, the descriptor for disease *d*(*i*) can be defined as follows:

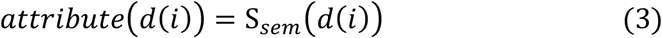

where *S_sem_*(*d*(*i*)) is a vector consisting of a collection of semantic similarities between disease *d*(*i*) and all diseases. Descriptors corresponding to 877 diseases in HMDD v3.0 were used to construct attribute information of the disease.

##### sequence descriptor of miRNA

The properties of the miRNA are represented by sequence information. For the sake of simplicity, *k*-mer is used to convert the sequence into a numerical vector, where *k* represents the length of the segmented subsequence [22]. For example, the 3-mer miRNA sequence can be expressed as AAA, UAA, … UUU. In this experiment, we use 3-mer to segment the sequence and use the frequency of the normalized 64 subsequences as the sequence descriptor *attribute* (*m*(*j*), where *m*(*j*) is the *j*th miRNA.

#### 2.2.2 Node representation

In order to effectively represent the relationship between each node and other nodes in the entire biological network, a network embedding method named Node2Vec is utilized in this study [23]. Node2Vec method is based on the sampling node neighborhood strategy of random walk, and optimizes the neighborhood preserving likelihood objective by the Skip-gram model to obtain the network embedded representation of the node. The method simulates a random walk of each node, wherein the */*-th node *c*(*i*) in the walk can be described as follows:

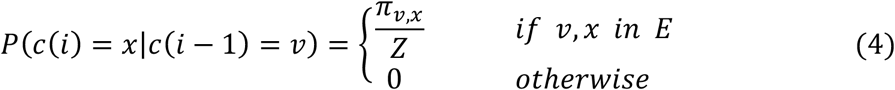

where *Z* is the normalization constant and *π_v,x_* is defined as the unstandardized transition probability of nodes *v* and *x*:

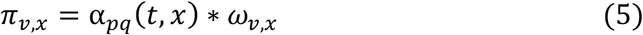

where *ω_v,x_* is the weight of the edges *v* and *x*, and the unweighted graph used in this experiment is set to 1. α_*pq*_ (*t, x*) is used to adjust the search process, interpolating between BFS and DFS. It is defined as follows:

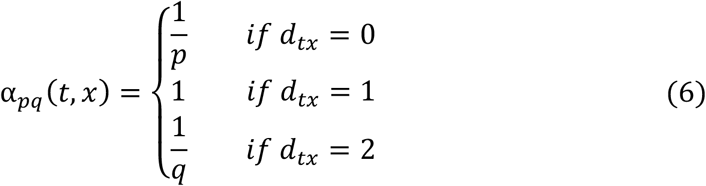

Where *d_tx_* is the shortest distance between node *t* and node *x. p* and *q* are the return parameter and the In-out parameter, respectively. And, the default parameters are used in this experiment. The specific details are shown in Figure 2. By learning the low-dimensional potential representation of nodes in the entire biological network, each node can describe its network relationship through a 128-dimensional vector *manner* (*node* (*i*)). *node*(*i*) is the *i*-th node in the network.

**Figure 2.**
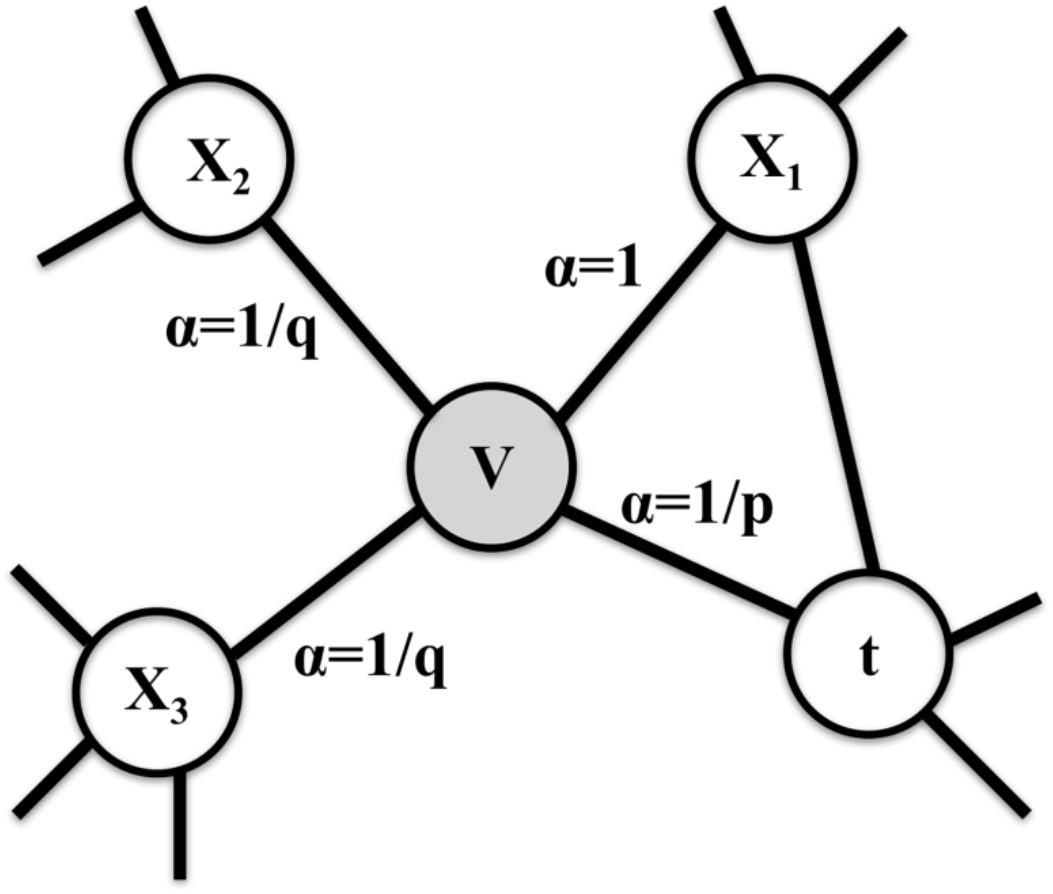
The 2nd-order biased random walks procedure in Node2Vec.

#### 2.2.3 Stacked autoencoder

Information from multiple sources is integrated in the proposed model, including information on the characteristics of diseases and miRNAs, as well as network representations of diseases and miRNAs. This operation allows the feature to contain more information, however, due to the different scope and size of the data from different sources, the model will be overly complex and prone to overfitting. Therefore, Stacked Autoencoder is adopted to obtain the appropriate subspace from the original feature space, which can effectively prevent over-fitting and improve the generalization performance of the model. The encoder that encodes the input *X* as a hidden representation *Y* is defined as follows:

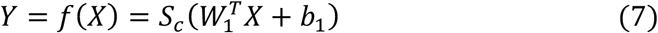

The decoder that maps the hidden representation *Y* to the approximate output *Z* is defined as follows:

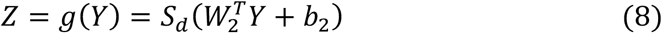

Where *S_c_* and *S_d_* are the activation functions. *W*_1_ and *W*_2_ are relational parameters. *b*_1_ and *b*_1_ are bias parameters.

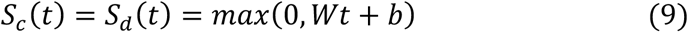

#### 2.2.4 Method overview

In this study, a new predictor called iMDA-BN was constructed to predict potential associations between miRNAs and diseases. The iMDA-BN is roughly divided into three parts, as shown in Figure 3. Firstly, node attribute, miRNA-based high-throughput data information and disease semantic information are used to construct miRNA sequence descriptors and disease semantic descriptors, respectively. Secondly, edge embedding, the network representation learning based on Biological Network calculates the node representation of each miRNA and disease. Finally, training random forest models to calculate miRNA-disease association scores. Next, the details in the experiment are described in detail.

**Figure 3.**
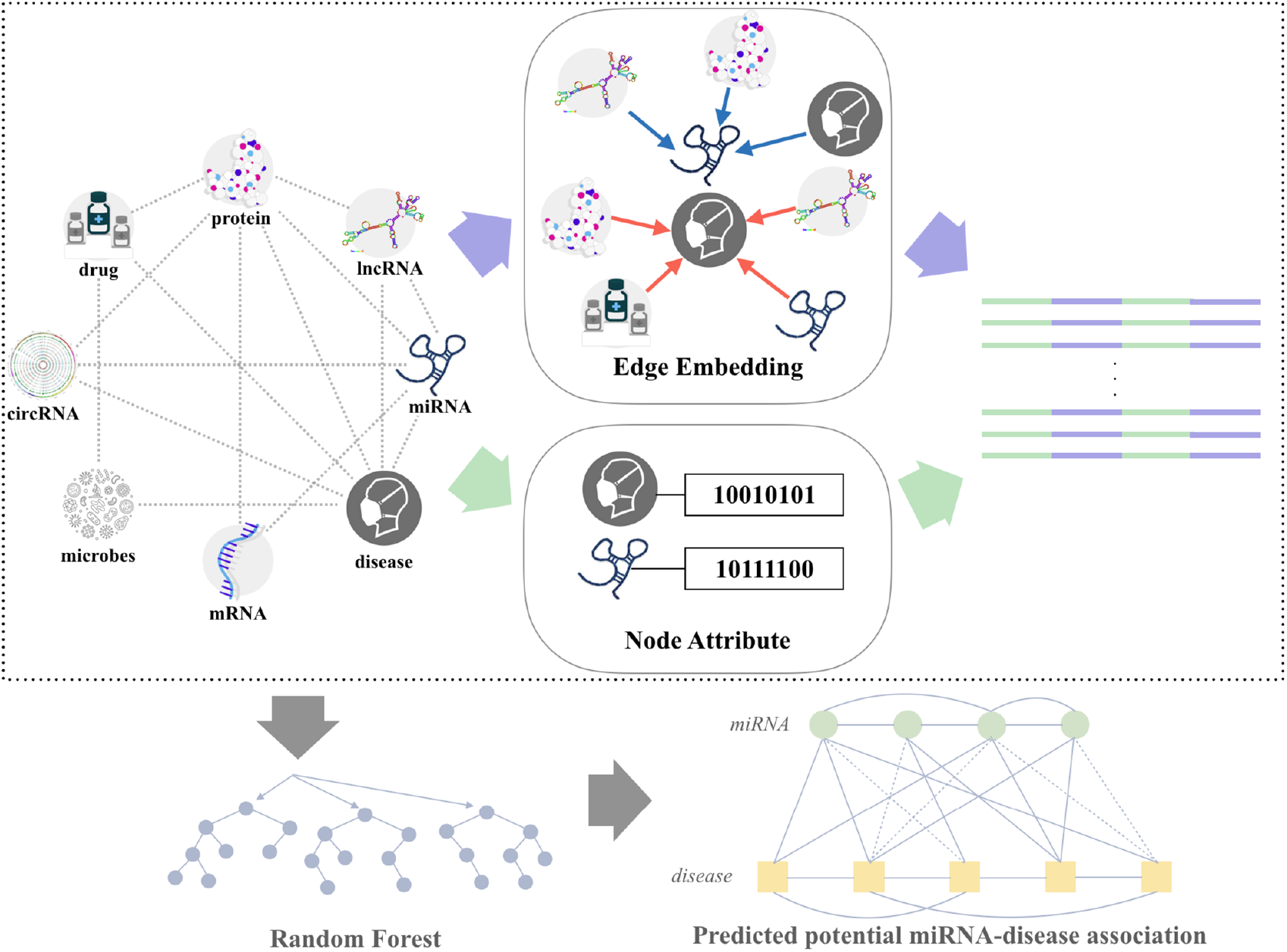
The framework of iMDA-BN.

##### The choice of positive and negative samples

Specifically, the 16427 miRNA-disease associations provided in HMDD are utilized as positive samples. The downsampling method was used to construct negative samples by randomly extracting the same number of associations from positive samples from 773,750 unconfirmed miRNA-disease associations. Although there may be potential associations, the number of negative samples we selected was only 16427 ÷ (901 × 877) ≈ 2.08% of the total number of samples. Therefore, there is a low probability of potential associations that can be ignored.

##### The construction of the final feature descriptor

As shown in Figure 3, the final feature descriptor of size 256 × 32854 is represented by the miRNA sequence descriptor, the miRNA node, the disease semantic descriptor and the disease node representation. The final association descriptor *F* of disease *d*(*j*) and miRNA *m*(*i*) can be described as a 256-dimensional vector:

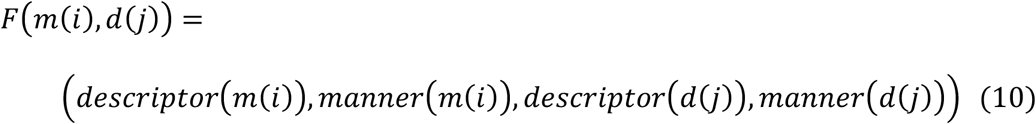

##### Prediction of miRNA-disease association by Random Forest

The final descriptor is used to train the random forest model and predict potential associations based on the trained model. In particular, the higher the prediction score, the more likely it is to be the candidate for potential associations.

## 3 Experimental Results

### 3.1 Performance of the new prediction method

In order to comprehensively evaluate the robustness and effectiveness of the iMDA-BN predictor, a 5-fold cross-validation was performed on the proposed model on the HMDD v3.0 dataset. It is divided into three steps: 1) The 32,854 associations used in this experiment were divided into five approximately equal and disjoint subsets. 2) One of the subsets was selected as the test set to test the model performance, and the remaining four subsets were used as the training set to train model. 3) Repeat step 2 until that all subsets are selected as test sets. Thus, five sets of experimental results were obtained, and we reported them in Table 3 and Figure 4, respectively. The area under the curve (AUC) is the area of the graph surrounded by the receiver operating characteristic curve (ROC) where the ROC is an evaluation criterion. It can be seen from Table 3 that the average AUC of the iMDA-BN has reached 0.9145, and the standard deviation is only 0.32%, which indicates that the proposed predictor is robust. In addition, the accuracy (Acc.), sensitivity (Sen.), precision (Prec.), specificity (Spec.), Matthews correlation coefficient (MCC) and the area under precision-recall (AUPR) of the proposed model are 84.49%, 84.20%, 84.79%, 84.70%, 68.99% and 91.92%. From the results of this part of the experiment, the method we proposed is discriminative.

**Table 3.** 5-fold cross-validation results performed by proposed model on HMDD v3.0 dataset.

| **Test set** | **Acc.(%)** | **Sen.(%)** | **Prec. (%)** | **Spec.(%)** | **MCC(%)** | **AUC** | **AUPR** |
| --- | --- | --- | --- | --- | --- | --- | --- |
| 1 | 84.14 | 83.84 | 84.45 | 84.35 | 68.29 | 0.9100 | 0.9148 |
| 2 | 84.57 | 84.51 | 84.63 | 84.61 | 69.14 | 0.9133 | 0.9168 |
| 3 | 85.04 | 85.39 | 84.69 | 84.8 | 70.09 | 0.9185 | 0.9216 |
| 4 | 84.57 | 84.02 | 85.12 | 84.95 | 69.15 | 0.9159 | 0.9214 |
| 5 | 84.15 | 83.25 | 85.04 | 84.77 | 68.3 | 0.9150 | 0.9218 |
| Average | **84.49±0.37** | **84.20±0.80** | **84.79±0.28** | **84.70±0.23** | **68.99±0.75** | **0.9145±0.0032** | **0.9192±0.0032** |

**Figure 4.**
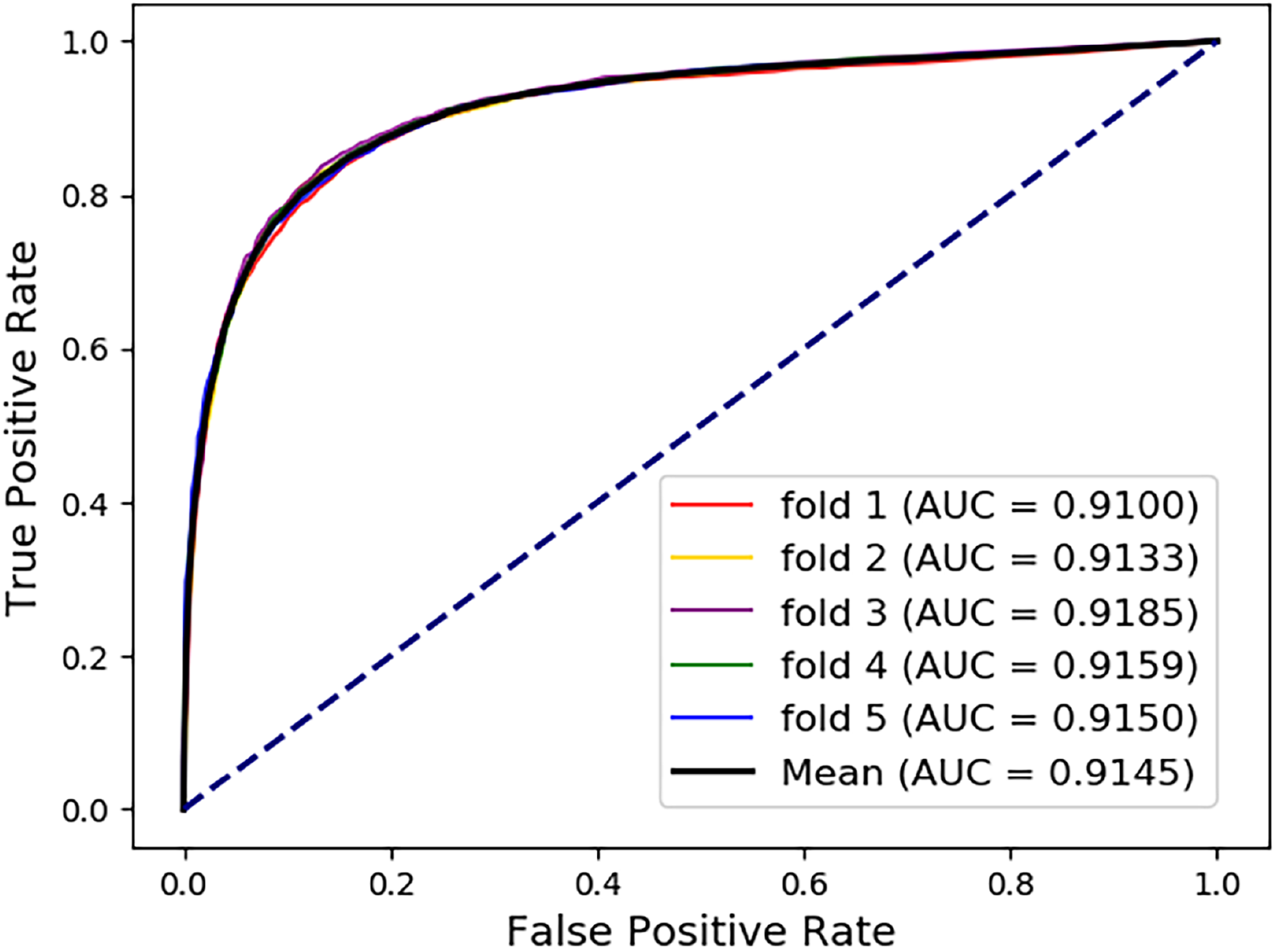
ROC curves performed by iMDA-BN.

**Figure 5.**
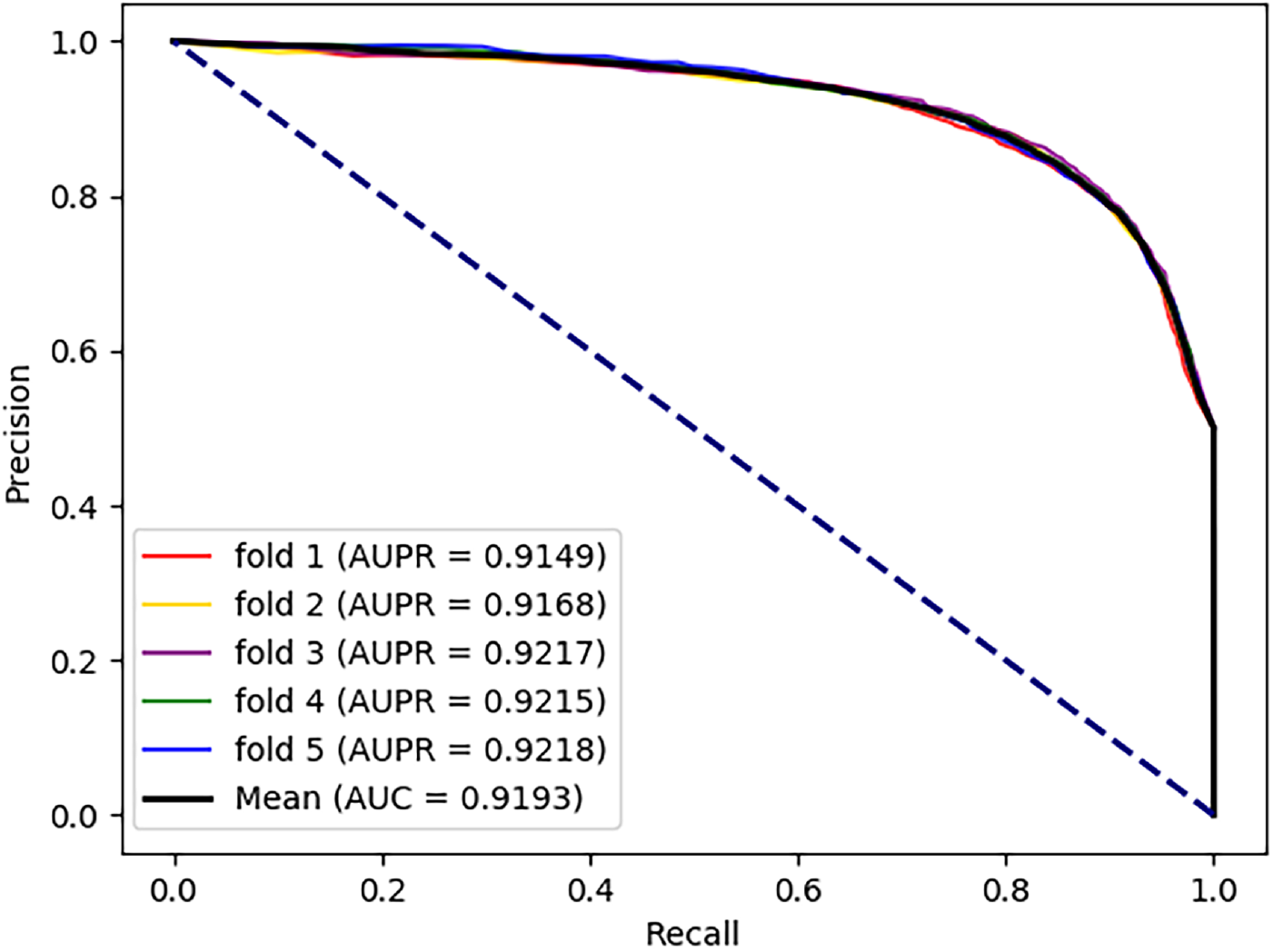
PR curves performed by iMDA-BN.

### 3.2 Compare different strategies to generate feature descriptors

Real-world networks, such as the biological network, are composed of nodes and edges, each associated with an essential attribute. In this method, the proposed feature descriptor *F*(*m*(*i*), *d*(*j*))is composed of the node attribute information *descriptor* and the node representation information *manner* that retains network structure information. In order to verify the reliability of the descriptors, we compare the three methods we implemented ourselves, using different descriptors in this experiment. Details are as follows.

- Descriptor “iMDA-BN (attribute)”: It consists of disease and miRNA attribute information, which can be described as *F_attribute_*(*m*(*i*), *d*(*j*), = *attribute* (m(i), d(j))
- Descriptor “iMDA-BN (manner)”: It consists of disease and miRNA node representation information, which can be described as *F_manner_*(*m*(*i*), *d*(*j*), = (*manner*(*m*(*i*), *manner* (*d*(*j*))).
- Descriptor “iMDA-BN”: The proposed feature descriptor *F*(*m*(*i*), *d*(*j*)

The above three descriptors all utilize the same random forest classifier, Medical Subject Headings and miRNA sequence information. Table 4 shows the scores of the three descriptors in the seven evaluation criteria including accuracy (Acc.), sensitivity (Sen.), specificity (Spec.), precision (Prec.) and Matthews correlation coefficient (MCC), AUC and AUPR. It can be seen that iMDA-BN outperforms other descriptors in the seven evaluation indicators, especially the AUC and MCC that measure the overall performance of the model. This suggests that multi-source knowledge that combines miRNA and disease attribute information with its node representation information in the network can describe the association between miRNA and disease from a more macro perspective, thereby characterizing the deeper meaning of multisource data.

**Table 4.** The comparison of different types of feature descriptors.

| Metric | Descriptor Comparison |  |  |
| --- | --- | --- | --- |
|  | iMDA-BN(Attribute) | iMDA-BN(Manner) | iMDA-BN |
| Acc.(%) | 80.96±0.53 | 84.12±0.28 | **84.49±0.37** |
| Sen.(%) | 83.79±1.12 | 83.85±0.79 | **84.20±0.80** |
| Prec.(%) | 78.14±0.35 | 84.40±0.29 | **84.79±0.28** |
| Spec.(%) | 79.30±0.28 | 84.31±0.15 | **84.70±0.23** |
| MCC(%) | 62.02±1.11 | 68.25±0.56 | **68.99±0.75** |
| AUC | 0.8765±0.0039 | 0.9106±0.0034 | **0.9145±0.0032** |
| AUPR | 0.8719±0.0036 | 0.9143±0.0031 | **0.9188±0.0030** |

Furthermore, for miRNAs and diseases that are not in the network, their characteristics can be characterized by combining their attribute information and setting the manner part to 0. The performance of iMAN-BN (attribute) is verified in Figure 6, indicating that the attribute information has considerable recognition. Therefore, in this way, we solve the problem of association prediction between miRNAs and diseases that are not in the network.

**Figure 6.**
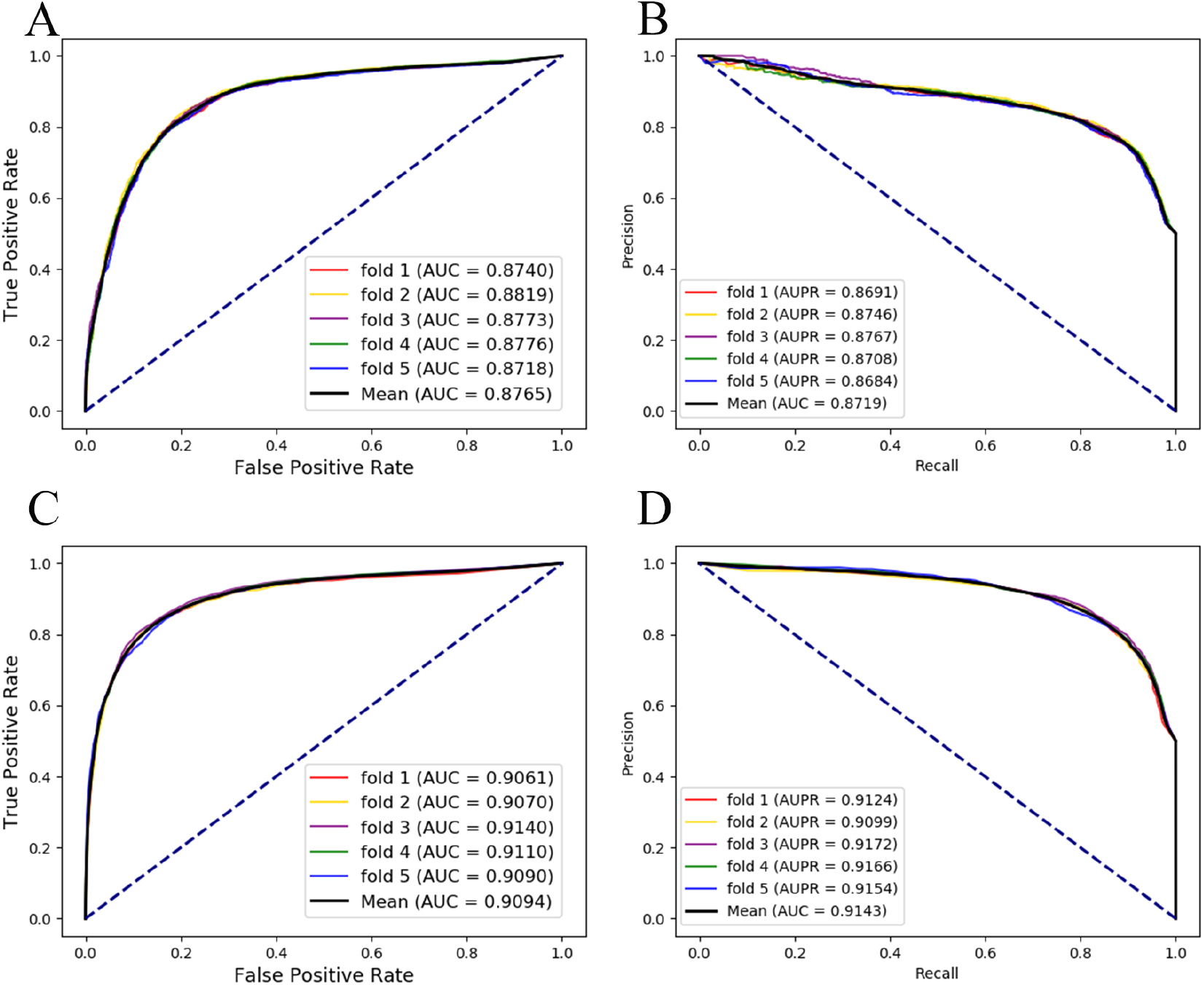
ROC and PR curves performed by iMDA-BN (Attribute) and iMDA-BN (manner). (A) ROC curves performed by iMDA-BN(Attribute). (B) PR curves performed by iMDA-BN (Attribute). (C) ROC curves performed by iMDA-BN (manner). (D) PR curves performed by iMDA-BN (manner).

### 3.3 Comparison with highly related methods

In recent years, many predictors have been proposed for the potential association between miRNAs and diseases. We compare the performance of the iMDA-BN with 7 state-of-the-art methods, including Shi’s, BNPMDA, LMTRDA, HGIMDA, BRWHNHA, KBMF-MDI and KBMF-MDI. Table 5 not only lists the performance of the various methods, but also shows the prior knowledge of building the associated network and the attribute information of the nodes. From the results, the iMDA-BN is superior to other methods using less than four associations on AUC and is 4.9% higher than the average. It is shown that the node representation information based on the biological network can improve the effect of predicting associations between miRNA and disease. In addition, the proposed method has an improvement of 2.48% and 15.65% in performance compared to methods that do not use attribute information such as MDA-CNN and Shi’s, which means that attribute information can also improve prediction performance. Furthermore, in Table 5, a protein-protein interaction (PPI) network of human genes was used as a gene network, as in previous studies, since miRNAs affect disease by regulating gene expression post-transcriptionally [24].

**Table 5.** The comparison with related models.

| Association | Methods | Attribute | AUC scores |
| --- | --- | --- | --- |
| PPI, MPI, DPI | Shi's<sup>1</sup> | N/A | 0.7580 |
|  | MDA-CNN<sup>2</sup> | N/A | 0.8897 |
| MDA | BNPMDA<sup>3</sup> | MeSH | 0.8980 |
|  | LMTRDA<sup>4</sup> | miRNA sequence, MeSH | 0.9054 |
|  | HGIMDA<sup>5</sup> | MeSH | 0.8781 |
|  | BRWHNHA<sup>6</sup> | MeSH | 0.8570 |
| MDA, DPI | KBMF-MDI<sup>7</sup> | miRNA sequence, MeSH, | 0.8725 |
| MDA, LDA, PPI, LMA,<br>LPI, DPI, MPI, PDI,<br>DDI | iMDA-BN | miRNA sequence, MeSH | 0.9145 |

<sup>1</sup>This method is reported in [6].
<sup>2</sup>This method is reported in [24].
<sup>3</sup>This method is reported in [9].
<sup>4</sup>This method is reported in [14].
<sup>5</sup>This method is reported in [38].
<sup>6</sup>This method is reported in [39].
<sup>7</sup>This method is reported in [40].

### 3.4 Case study

In this part of the experiment, the iMAN-BN’s the ability of predicting disease-associated miRNAs was validated by case studies of three common human diseases, assuming that prior knowledge are only associations in HMDD v3.0. Specifically, the training set is made up of all the associations in the final descriptor. At the same time, we used the associations that did not appear in HMDD for the three diseases as a test set. After iMAN-BN gave prediction scores to the test set, the top 50 miRNAs with the highest score for each disease were validated in the dbDEMC database and the miR2Disease database [16, 17]. Breast cancer is the most common female cancer in developed countries [25]. Its incidence increases rapidly with age, but its incidence decreases near the age of menopause [25]. Since some of the pathogenic factors of breast cancer are endogenous, this makes prevention very difficult. Recent studies have shown that mir-125b, mir-145, mir-21 and mir-155 in breast cancer tissues are significantly dysregulated compared to normal breast tissue [26]. In Table 6, we predicted potential breast neoplasms-associated miRNAs and verified the top 50 miRNAs with the highest scores, 45 of these miRNA-disease associations were confirmed. Colon cancer is the second most common cancer [27]. Since some colon cancer cells still cannot be eradicated by existing therapies, the study of the pathogenic principle has been a hotspot in biomedical research [27]. Studies have shown that the promoters of hsa-miR-9, hsa-miR-129 and hsa-miR-137 are abnormally hypermethylated in colon cancer cells [28]. In Table 7, we predicted potential Colon Neoplasms-associated miRNAs and verified the top 50 miRNAs with the highest scores, of which 49 miRNA-disease associations were confirmed. Lymphoma is a blood cancer that develops from lymphocytes and originates from lymphocytes [29]. In Table 8, we predicted potential Lymphoma-associated miRNAs and verified the top 50 miRNAs with the highest scores, 49 of these miRNA-disease associations were confirmed.

**Table 6.** Prediction of the top 50 predicted miRNAs associated with Breast Neoplasms.

| miRNA | dbDEMC | miR2D | miRNA | dbDEMC | miR2D |
| --- | --- | --- | --- | --- | --- |
| hsa-mir-181d-5p | confirmed | unconfirmed | hsa-mir-508-5p | confirmed | unconfirmed |
| hsa-mir-99b-5p | confirmed | unconfirmed | hsa-mir-154-5p | confirmed | unconfirmed |
| hsa-mir-330-5p | confirmed | unconfirmed | hsa-mir-581 | confirmed | unconfirmed |
| hsa-mir-28-5p | confirmed | unconfirmed | hsa-mir-501-5p | confirmed | unconfirmed |
| hsa-mir-361-5p | confirmed | unconfirmed | hsa-mir-323a-5p | confirmed | unconfirmed |
| hsa-mir-371a-5p | confirmed | unconfirmed | hsa-mir-628-5p | confirmed | unconfirmed |
| hsa-mir-885-5p | confirmed | unconfirmed | hsa-mir-612 | unconfirmed | unconfirmed |
| hsa-mir-455-5p | confirmed | unconfirmed | hsa-mir-490-5p | confirmed | unconfirmed |
| hsa-mir-651-5p | confirmed | unconfirmed | hsa-mir-188-5p | confirmed | unconfirmed |
| hsa-mir-1271-5p | confirmed | unconfirmed | hsa-mir-1299 | confirmed | unconfirmed |
| hsa-mir-504-5p | confirmed | unconfirmed | hsa-mir-95-5p | confirmed | unconfirmed |
| hsa-mir-876-5p | confirmed | unconfirmed | hsa-mir-1296-5p | confirmed | unconfirmed |
| hsa-mir-454-5p | confirmed | unconfirmed | hsa-mir-582-5p | confirmed | unconfirmed |
| hsa-mir-532-5p | confirmed | unconfirmed | hsa-mir-512-5p | confirmed | unconfirmed |
| hsa-mir-1297 | confirmed | unconfirmed | hsa-mir-1303 | confirmed | unconfirmed |
| hsa-mir-449b-5p | confirmed | unconfirmed | hsa-mir-323b-5p | confirmed | unconfirmed |
| hsa-mir-433-5p | confirmed | unconfirmed | hsa-mir-889-5p | confirmed | unconfirmed |
| hsa-mir-544a | confirmed | unconfirmed | hsa-mir-1184 | confirmed | unconfirmed |
| hsa-mir-136-5p | confirmed | confirmed | hsa-mir-500a-5p | confirmed | unconfirmed |
| hsa-mir-23c | unconfirmed | unconfirmed | hsa-mir-217-5p | confirmed | unconfirmed |
| hsa-mir-761 | unconfirmed | unconfirmed | hsa-mir-518e-5p | confirmed | unconfirmed |
| hsa-mir-4500 | unconfirmed | unconfirmed | hsa-mir-376b-5p | confirmed | unconfirmed |
| hsa-mir-346 | confirmed | unconfirmed | hsa-mir-186-5p | confirmed | unconfirmed |
| hsa-mir-216a-5p | confirmed | unconfirmed | hsa-mir-498-5p | confirmed | unconfirmed |
| hsa-mir-382-5p | confirmed | unconfirmed | hsa-mir-764 | unconfirmed | unconfirmed |

**Table 7.** Prediction of the top 50 predicted miRNAs associated with Colon Neoplasms.

| miRNA | dbDEMC | miR2D | miRNA | dbDEMC | miR2D |
| --- | --- | --- | --- | --- | --- |
| hsa-mir-16-5p | confirmed | unconfirmed | hsa-mir-505-5p | confirmed | unconfirmed |
| hsa-mir-29c-5p | confirmed | unconfirmed | hsa-mir-495-5p | confirmed | unconfirmed |
| hsa-mir-423-5p | confirmed | unconfirmed | hsa-mir-122-5p | confirmed | unconfirmed |
| hsa-mir-146b-5p | confirmed | unconfirmed | hsa-mir-34b-5p | confirmed | confirmed |
| hsa-mir-193a-5p | confirmed | unconfirmed | hsa-mir-7-5p | confirmed | confirmed |
| hsa-mir-98-5p | confirmed | unconfirmed | hsa-mir-370-5p | confirmed | unconfirmed |
| hsa-mir-124-5p | confirmed | confirmed | hsa-mir-34c-5p | confirmed | confirmed |
| hsa-mir-9-5p | confirmed | confirmed | hsa-mir-134-5p | confirmed | unconfirmed |
| hsa-mir-130b-5p | confirmed | confirmed | hsa-mir-491-5p | confirmed | unconfirmed |
| hsa-mir-128-3p | confirmed | confirmed | hsa-mir-212-5p | confirmed | unconfirmed |
| hsa-mir-199a-5p | confirmed | unconfirmed | hsa-mir-149-5p | confirmed | unconfirmed |
| hsa-mir-362-5p | unconfirmed | unconfirmed | hsa-mir-129-5p | confirmed | confirmed |
| hsa-mir-372-5p | confirmed | confirmed | hsa-mir-181a-2-3p | confirmed | confirmed |
| hsa-mir-27b-5p | confirmed | confirmed | hsa-mir-99b-5p | confirmed | unconfirmed |
| hsa-mir-494-5p | confirmed | unconfirmed | hsa-mir-144-5p | confirmed | unconfirmed |
| hsa-mir-139-5p | confirmed | confirmed | hsa-mir-182-5p | confirmed | confirmed |
| hsa-mir-92b-5p | confirmed | unconfirmed | hsa-mir-99a-5p | confirmed | unconfirmed |
| hsa-mir-10a-5p | confirmed | confirmed | hsa-mir-373-5p | confirmed | unconfirmed |
| hsa-mir-92a-2-5p | confirmed | unconfirmed | hsa-mir-29b-2-5p | confirmed | confirmed |
| hsa-mir-199b-5p | confirmed | unconfirmed | hsa-mir-20b-5p | confirmed | unconfirmed |
| hsa-mir-214-5p | confirmed | unconfirmed | hsa-mir-320a-5p | confirmed | unconfirmed |
| hsa-mir-217-5p | confirmed | unconfirmed | hsa-mir-28-5p | confirmed | unconfirmed |
| hsa-mir-590-5p | confirmed | unconfirmed | hsa-mir-26a-2-3p | confirmed | confirmed |
| hsa-mir-342-5p | confirmed | confirmed | hsa-mir-100-5p | confirmed | unconfirmed |
| hsa-mir-421 | confirmed | unconfirmed | hsa-mir-302c-5p | confirmed | unconfirmed |

**Table 8.** Prediction of the top 50 predicted miRNAs associated with Lymphomas.

| miRNA | dbDEMC | miR2D | miRNA | dbDEMC | miR2D |
| --- | --- | --- | --- | --- | --- |
| hsa-mir-145-5p | confirmed | confirmed | hsa-mir-876-5p | confirmed | unconfirmed |
| hsa-let-7b-5p | confirmed | unconfirmed | hsa-let-7e-5p | confirmed | confirmed |
| hsa-let-7a-5p | confirmed | confirmed | hsa-mir-15b-5p | confirmed | unconfirmed |
| hsa-mir-182-5p | confirmed | unconfirmed | hsa-mir-199b-5p | confirmed | unconfirmed |
| hsa-mir-34a-5p | confirmed | unconfirmed | hsa-mir-106b-5p | confirmed | unconfirmed |
| hsa-mir-107 | confirmed | unconfirmed | hsa-mir-192-5p | confirmed | unconfirmed |
| hsa-mir-424-5p | confirmed | unconfirmed | hsa-mir-106a-5p | confirmed | confirmed |
| hsa-mir-98-5p | confirmed | unconfirmed | hsa-mir-146b-5p | confirmed | unconfirmed |
| hsa-mir-195-5p | confirmed | unconfirmed | hsa-mir-33b-5p | confirmed | unconfirmed |
| hsa-mir-181b-5p | confirmed | unconfirmed | hsa-mir-196a-5p | confirmed | confirmed |
| hsa-mir-9-5p | confirmed | confirmed | hsa-mir-33a-5p | confirmed | unconfirmed |
| hsa-mir-218-5p | confirmed | unconfirmed | hsa-mir-125b-1-3p | unconfirmed | unconfirmed |
| hsa-mir-335-5p | confirmed | unconfirmed | hsa-mir-181d-5p | confirmed | unconfirmed |
| hsa-mir-196b-5p | confirmed | unconfirmed | hsa-mir-346 | confirmed | unconfirmed |
| hsa-mir-138-5p | confirmed | unconfirmed | hsa-mir-100-5p | confirmed | unconfirmed |
| hsa-mir-29a-5p | confirmed | unconfirmed | hsa-mir-590-5p | confirmed | unconfirmed |
| hsa-let-7g-5p | confirmed | unconfirmed | hsa-mir-361-5p | confirmed | unconfirmed |
| hsa-mir-30b-5p | confirmed | confirmed | hsa-mir-421 | confirmed | unconfirmed |
| hsa-mir-503-5p | confirmed | unconfirmed | hsa-mir-320b | confirmed | unconfirmed |
| hsa-mir-24-3p | confirmed | unconfirmed | hsa-mir-7-5p | confirmed | unconfirmed |
| hsa-mir-125b-5p | confirmed | unconfirmed | hsa-mir-149-5p | confirmed | confirmed |
| hsa-mir-374a-5p | confirmed | unconfirmed | hsa-mir-32-5p | confirmed | unconfirmed |
| hsa-mir-326 | confirmed | unconfirmed | hsa-mir-216b-5p | confirmed | unconfirmed |
| hsa-mir-134-5p | confirmed | unconfirmed | hsa-mir-129-5p | confirmed | unconfirmed |
| hsa-mir-136-5p | confirmed | unconfirmed | hsa-let-7i-5p | confirmed | unconfirmed |

## 4 Conclusion

With the development of bioinformatics, more and more predictors of potential associations have been proposed, and these methods have greatly promoted the development of biomedicine. However, they focus only on the association network of research content, and methods based on the entire biological network are scarce. Therefore, it is necessary to develop a biological network-based computational method to identify the association between potential miRNAs and diseases. In this paper, we propose a novel computational method based on a complex biological network composed of nine associations called iMDA-BN to predict the potential association between potential miRNAs and disease. From the experimental results, it is better than other most advanced methods, and it can predict the association between miRNA and disease that does not exist in the network. In addition, we also demonstrated the excellent ability of iMDA-BN to predict potential associations through three case studies, and achieved 90%, 98% and 98% accuracy. The reliability of iMDA-BN can be achieved mainly for three reasons: I) It uses a new method to describe disease and miRNA characteristics which analyzes node representation information for disease and miRNA from the perspective of biological networks. II) It can predict unproven associations even if miRNAs and diseases do not appear in the biological network. III) Accurate description of miRNA characteristics from biological properties based on high-throughput sequence information.

### Key points

- It uses a new method to describe disease and miRNA characteristics which analyzes node representation information for disease and miRNA from the perspective of biological networks.
- It can predict unproven associations even if miRNAs and diseases do not appear in the biological network.
- Accurate description of miRNA characteristics from biological properties based on high-throughput sequence information.

## Author biographies

**Kai Zheng**, master, is currently a master from China University of Mining and Technology. His research interests include data mining, pattern recognition, recommender systems, machine learning, deep learning, intelligent information processing and its applications in bioinformatics.

**Zhu-Hong You**, PhD, is a professor of the Xinjiang Technical Institutes of Physics and Chemistry, Chinese Academy of Sciences. His research interests include disease and circular RNAs, network pharmacology, and machine learning.

**Lei Wang**, PhD, is an assistant research fellow of the Xinjiang Technical Institutes of Physics and Chemistry, Chinese Academy of Sciences. His research interests include disease and circular RNAs, network pharmacology, and machine learning.

## Acknowledgements

This work is supported is supported in part by Awardee of the NSFC Excellent Young Scholars Program, under Grants 61722212, in part by the National Science Foundation of China, under Grants 61873212, 61702444, 61572506, in part by the Pioneer Hundred Talents Program of Chinese Academy of Sciences, in part by the the West Light Foundation of The Chinese Academy of Sciences, under Grant 2018-XBQNXZ-B-008. The authors would like to thank all anonymous reviewers for their constructive advices.

